# Identification and Characterization of CtUGT3 as the Key Player of Astragalin Biosynthesis in Safflower

**DOI:** 10.1101/2023.06.22.546132

**Authors:** Chaoxiang Ren, Ziqing Xi, Bin Xian, Chao Chen, Xulong Huang, Huajuan Jiang, Jiang Chen, Cheng Peng, Jin Pei

**Author notes:** Corresponding Authors: Jiang Chen, Cheng Peng Jin Pei. These authors contributed equally to this work (Chaoxiang Ren, Ziqing Xi).

## Abstract

Safflower (*Carthamus tinctorius* L.) flowers are used as a traditional Chinese medicine for a long history. Flavonoids are the main bioactive components in safflower flowers, and most of them exist in the form of flavonoid glycosides. Only few glycosyltransferases have been identified in safflower. To reveal the uridine diphosphate glycosyltransferase (UGT) involved in flavonoid glycosides biosynthesis in safflower, a metabolomics and transcriptome analysis was performed by using the flowers under different light qualities treatments. Three differentially expressed *UGT* genes were screened, and their expressions were significantly related with concentrations of 9 flavonoid *O*-glycosides. Safflower corolla protoplasts were further used to confirm flavonoid *O*-glycosylation ability of UGT candidates. The astragalin (kaempferol 3-*O*-glucoside) content was only significantly increased when CtUGT3 was overexpressed in protoplasts. The biochemical properties and kinetic parameters of CtUGT3 were determined. CtUGT3 also showed flavonoid 3-OH and 7-OH glycosylation activities *in vitro*. Molecular modeling and site-directed mutagenesis revealed that E384 and S276 were critical catalytic residues for the 3-OH glycosylation of CtUGT3. These results demonstrate that CtUGT3 has a flavonoid 3-OH glycosylation function and is involved in the biosynthesis of astragalin in safflower. This study provides insights into the catalytic mechanisms of flavonoid *O*-glycosyltransferases, and makes a reference for flavonoid biosynthesis genes research in medicinal plants.

## Introduction

Safflower (*Carthamus tinctorius* L.) is an annual Asteraceae species and the only cultivar of the genus *Carthamus*. Safflower is a multipurpose economic crop that is distributed worldwide. Its seeds are used for the production of edible oil, and the yellow and red pigments, which are widely used in food, medicines, cosmetics, and other products, are extracted from its flowers (Azami et al. 2019). In East Asian countries such as China, Japan, Korea, and Thailand, safflower flowers have been used in traditional Chinese medicine for a long history. Safflower flowers are used to treat various diseases, such as cerebrovascular, cardiovascular, and gynecological diseases (Delshad et al. 2018). Flavonoids are the main bioactive components in safflower flowers (Yue et al. 2013; Zhang et al. 2016; Xue et al. 2021) and exhibit several pharmacological properties such as anti-inflammatory, anti-cancer, and anti-ischemic effects on the heart and brain, as well as protective effects on the liver and lungs (Xian et al. 2022).

However, most flavonoids in plants exist as glycosides, and the most abundant flavonoid glycosides are *O*-glycosides and *C*-glycosides (Veitch and Grayer 2011; Xiao 2017). Glycosylation is a prominent modification that is usually the last step in the biosynthesis of natural compounds, in coordination with hydroxylation, acylation, and methylation. Kaempferol is one of the most common flavonoid sugar acceptors and always binds to a single sugar or multiple sugars at the 3, 6, or 7 positions to form *O*-glycosides, di-*O*-glycosides, or tri-*O*-glycoside (Fan et al. 2009; Xie et al. 2016). Kaempferol 3-*O*-glucoside (astragalin) is one of the most abundant bioactive compounds found in safflower flowers. Astragalin is found in many medicinal plants with various biological activities, such as anti-tumor, anti-inflammatory, anti-allergic, anti-oxidant, and anti-bacterial activities (Riaz et al. 2018).

The basic metabolic pathway of flavonoid biosynthesis is well known, particularly in model plants (Saito et al. 2013). In safflower flowers, many functional genes involved in flavonoid biosynthesis have been cloned and analyzed, including *CHS*, *CHI*, *FLS*, and *F3H* (Shinozaki et al. 2016; Tu et al. 2016; Guo et al. 2019; Tu et al. 2019). In our previous study, genes involved in flavonoid biosynthesis were screened and cloned based on transcriptome and expression analysis (Chen et al. 2018; Ren et al. 2019). However, the flavonoid glycosyltransferases (GTs) in safflower are largely unknown. Several studies have been published on the GTs in different plants, including *O*-GTs (Nawade et al. 2020; Huang et al. 2021; Dai et al. 2022), *C*-GTs, and di-*C*-GTs (Liu et al. 2020; Ren et al. 2020c; Zhang et al. 2020; Feng et al. 2021; Huang et al. 2022). However, to date, only two GTs have been identified in safflower, and their functions have only been confirmed *in vitro* (Xie et al. 2014; Guo et al. 2016). It is necessary to screen and identify new glycosyltransferases in *Carthamus tinctorius* because safflower contains abundant bioactive flavonoid glycosides.

On the other hand, researchers found that temperature, precipitation, light intensity, and drought stress affected the growth, distribution, and secondary metabolic accumulation of safflower (Wei et al. 2018; Ren et al. 2020a; Wei et al. 2020). Light is an important environmental factor that regulates plant growth and developmental cycles (de Wit et al. 2016). Sunlight consists of ultraviolet (UV), visible (light), and infra-red light. Generally, blue, green, and red light are the major components that positively influence the yield, growth, and nutrient quality of plants. Regarding secondary metabolism in plants, several studies have reported that red, blue, and UV light affect biosynthesis and accumulation (Henry-Kirk et al. 2018; Liu et al. 2018; Zhang et al. 2018; Zhang et al. 2019). However, in specific situations, other light colors are also beneficial. Therefore, different light quality treatments offer an effective method for screening genes involved in flavonoid glycosylation.

In this study, to reveal the uridine diphosphate glycosyltransferase (UGT) involved in flavonoid glycoside biosynthesis in safflower flowers, we performed metabolomics and transcriptome analyses by using flowers under different light quality treatments. Three UGT candidates related to flavonoid glycosides content were screened. Safflower corolla protoplasts were used to confirm the flavonoid *O*-glycosylation ability of UGT candidates. The enzyme assays were performed *in vitro*. The biochemical properties and kinetic parameters of CtUGT3 were also determined. Molecular modeling and site-directed mutagenesis were used to further understand the catalytic mechanism of CtUGT3. Our results confirm that CtUGT3 exhibits *O*-GT activity and is involved in astragalin biosynthesis in safflower.

## Results

### Flavonoid metabolites and transcript analysis among safflower samples under different light quality treatments

The flavonoid metabolites of the four groups of safflowers treated with blue light (Bl), far-red light (FR), red light (Re), and UV-B (UV) were compared to those of safflowers grown under white light (Wh) and dark conditions (Dr), respectively. A total of 114 flavonoid metabolites were detected (Supplemental Table S1), 32 of which were differentially metabolized (Supplemental Table S2). Compared with that observed under white light, safflower exhibited the greatest differences in flavonoid metabolites under far-red light, UV-B, and dark conditions. Compared with those grown under dark conditions, the flavonoid metabolites of safflower grown under far-red light and UV-B exhibited the greatest differences (Figure 1A). Thirty-two differentially metabolized flavonoids belonged to 7 different secondary classification components, including flavanol, flavonoid carboglycoside, isoflavone, flavone, flavonol, orange flavonoid, and dihydroflavonoid. Among them, flavonols and flavones had the most differential metabolites with 11 and 9 species, respectively (Figure 1B). In addition, 18 of the 32 flavonoids were glycosides, including 15 *O*-glycosides and 3 *C*-glycosides. Most of the aglycones of these glycosides had flavonol or flavone structures.

**Figure 1.**
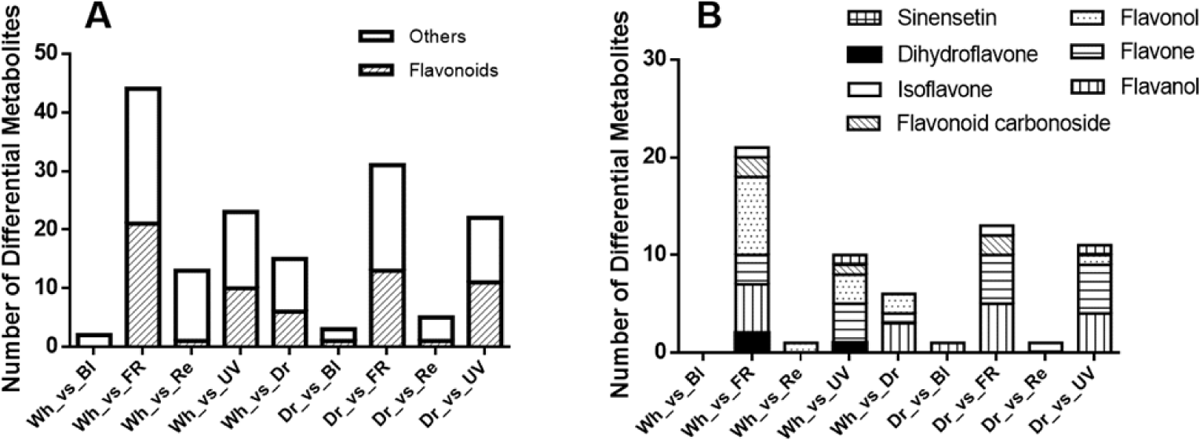
Different flavonoid metabolites of safflower cultivated under different light quality conditions. The x-axis represents the comparisons of different light conditions, and the Y-axis represents the number of different metabolites.

Through transcript analysis, a total of 121 Gb of clean data was obtained. The Q30 value of each sample was greater than 91%. The clean reads quality metrics are listed in Supplemental Table S3. More than 95% of clean reads were mapped to the reference genome (Supplemental Table S4). Based on gene expression levels, 1269 differentially expressed genes (DEGs) were identified (Supplemental Table S5). Compared with white light or dark conditions, DEGs in the FR group were the most abundant, followed by those in the UV group (Supplemental Table S6). Eight DEGs were annotated to the flavonoid biosynthesis pathway, including two *PAL*, *CHS*, *HCT*, *FLS* and three *UGT*, named as *CtPAL1*, *CtPAL2*, *CtCHS*, *CtHCT*, *CtFLS*, *CtUGT1*, *CtUGT2* and *CtUGT3* respectively. Eight DEGs were verified by RT-qPCR. The RT-qPCR results were consistent with those of RNA-Seq (Supplemental Figure S1), indicating that the sequencing data were accurate. Expression of *CtUGT1* increased under blue and UV light and decreased under FR light. Expression of *CtUGT2* increased under blue and red light and decreased in the dark. Expression of *CtUGT3* increased under blue light and decreased under FR and dark conditions (Figure 2). Flavonoid glycosides that were significantly related to the three differentially expressed *UGT* genes were screened. Finally, the contents of 9 flavonoid glycosides were significantly correlated with the expression of *CtUGT1*, *CtUGT2*, and *CtUGT3*, mainly 3-*O* glycosides and 7-*O* glycosides of flavonol (Table 1).

**Figure 2.**
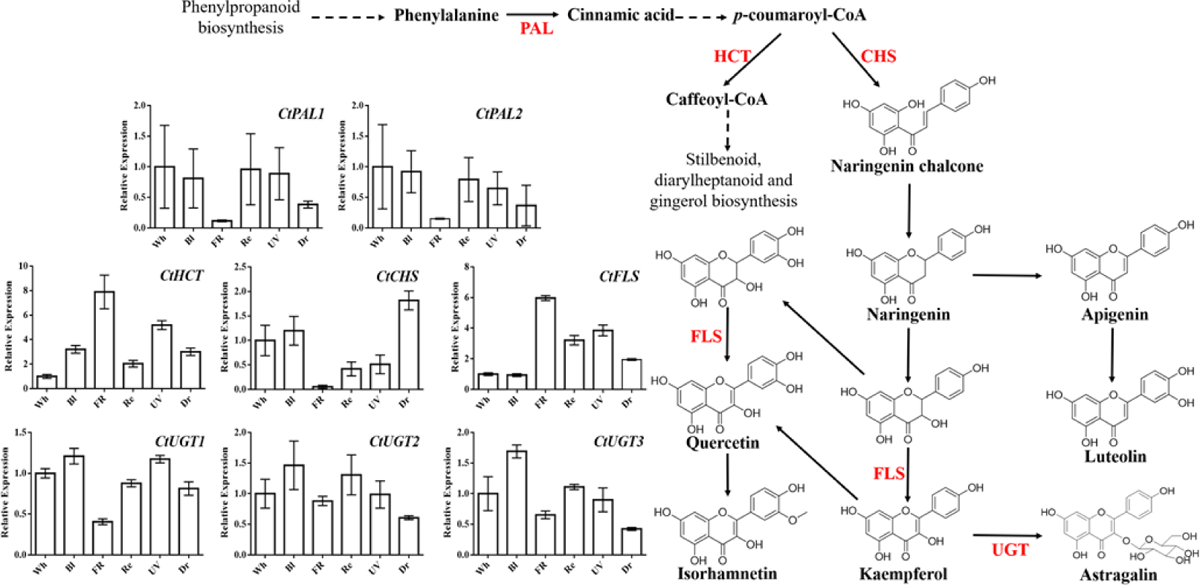
Relative expressions of 8 differential expressed genes related with flavonoid biosynthesis pathway.

**Table 1.** Significantly correlated differentially expressed genes and metabolites

| Genes | Compounds | Type | Correlation |
| --- | --- | --- | --- |
| *CtUGT1* | Diosmetin 7-*O*-glucuronide | Flavone | Positive |
|  | Kaempferol 7-*O*-glucosdie | Flavonol | Positive |
|  | Trifolin (Kaempferol 3-*O*-galactoside) | Flavonol | Positive |
|  | Astragalin (Kaempferol 3-*O*-glucosdie) | Flavonol | Positive |
|  | Luteolin 7-*O*-glucoside | Flavone | Positive |
|  | Quercetin 7-*O*-glucoside | Flavonol | Positive |
|  | Isorhamnetin 3-*O*-rutinoside | Flavonol | Negative |
|  | Quercetin 3-*O*-robinobioside | Flavonol | Negative |
| *CtUGT2* | Luteolin 7-*O*-glucoside | Flavone | Positive |
|  | Isorhamnetin 3-*O*-glucuronide | Flavonol | Positive |
| *CtUGT3* | Kaempferol 7-*O*-glucosdie | Flavonol | Positive |
|  | Trifolin (Kaempferol 3-*O*-galactoside) | Flavonol | Positive |
|  | Astragalin (Kaempferol 3-*O*-glucosdie) | Flavonol | Positive |
|  | Luteolin 7-*O*-glucoside | Flavone | Positive |
|  | Quercetin 7-*O*-glucoside | Flavonol | Positive |
|  | Isorhamnetin 3-*O*-glucuronide | Flavonol | Positive |
|  | Isorhamnetin 3-*O*-rutinoside | Flavonol | Negative |
|  | Quercetin 3-*O*-robinobioside | Flavonol | Negative |

### Cloning and bioinformatics analysis of UGT candidates

The full-length coding sequences of the three differentially expressed *UGTs* were cloned at 1425, 1431, and 1425 bp, respectively (GenBank accession number: OQ354214, OQ354222, OQ354223). The predicted molecular weights, pIs, extinction coefficients and aliphatic indices are shown in Supplemental Table S7. Phylogenetic analysis revealed that the three UGT candidates were closely related to GT04F14, TcCGT, and UGT71E5. GT04F14 could catalyze the 7-OH group of isoflavones (He et al. 2011). TcCGT exhibited both *C*- and *O*-glycosylation activity toward several flavonoid substrates (He et al. 2019). UGT71E5 showed high catalytic efficiency in *O*-glycosylation with flavonoids (Xie et al. 2017). It was speculated that candidate CtUGTs might have functions similar to those of enzymes in the same clade (Figure 3).

**Figure 3.**
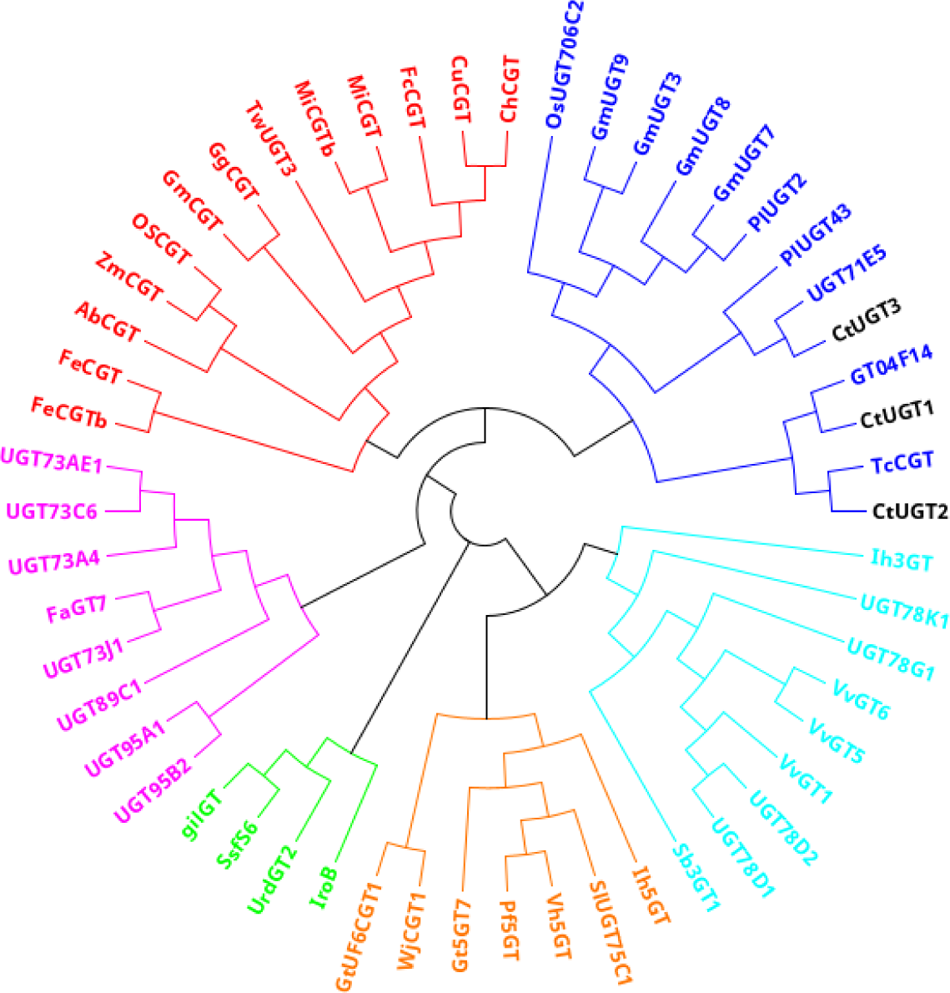
Phylogenetic analysis of UGTs.

### Transient expression of CtUGT3 in safflower corolla protoplast

Safflower flowers contain some special flavonoids such as chalcone *C*-glycosides represented by hydroxysafflor yellow A (HSYA). However, model plants commonly used for functional identification may not contain these components or their precursors. Therefore, the safflower corolla protoplast is ideal for verifying the functional genes involved in safflower flavonoid biosynthesis. Protoplasts were successfully isolated from the tender corollas of safflower flowers. The pA7-CaMV35S-CtUGT3-YFP vector was constructed and transformed into corolla protoplasts. Safflower corolla protoplasts contain light-yellow cells that emit green or yellow-green fluorescence at an excitation wavelength of around 480 ∼ 490 nm. Yellow fluorescent protein (YFP), which emit green, yellow, or red fluorescence at different wavelengths, was selected as the fluorescent marker protein. The DsRed (558 nm) excitation wavelength was selected for observation in fluorescence microscopy. Wild-type cells exhibited no fluorescence in the presence of DsRed, but cells that successfully transiently expressed the YFP tag showed red fluorescence. Therefore, detection of YFP under DsRed could satisfactorily distinguish the self-fluorescence of yellow safflower protoplasts from the transient expression of fluorescent markers.

In the protoplasts that were successfully transformed into *CtUGT1*, *CtUGT2*, and *CtUGT3* respectively, there were obvious red fluorescence under DsRed, and some cell’ colors changed from transparent or light yellow to deep yellow or even red (Figure 4). This result indicated that the transient expression of CtUGTs might change the accumulation of flavonoids in the protoplasts.

**Figure 4.**
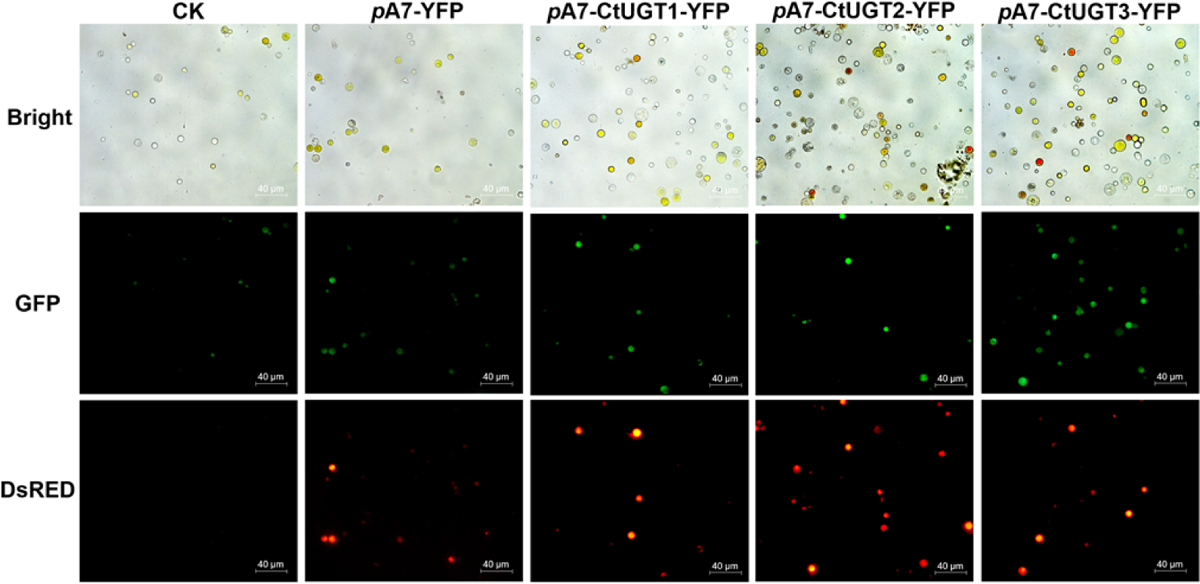
Transient expression of CtUGTs in safflower corolla protoplast. CK represents the untransfected group. *p*A7-YFP represents the group of transfected empty vector. *p*A7-CtUGTs-YFP represent the group of transfected vectors with CtUGTs genes. Bright represents the photographs was captured under bright light. GFP represents the photographs was captured under 488 nm. DsRED represents the photographs was captured under 558 nm.

The flavonoid glycosides in safflower protoplasts were determined using UPLC-QTOF-MS/MS. Six peaks were identified using commercial flavonoid glycoside standards: 6-hydroxyl kaempferol 3,6-*O*-diglucosyl-7-*O*-glucuronic acid (HKDG); 6-hydroxyl kaempferol 3,6,7-*O*-triglucoside (HKTG); HSYA; anhydrosafflor yellow B (AB); kaempferol 3-*O*-rutinoside (K3OR); and astragalin (kaempferol 3-*O*-glucoside) (Supplemental Figure S2-S7). It was found that the HKDG content decreased and the astragalin content was more than doubled when *CtUGT3* was overexpressed in protoplast cells (Figure 5). The concentrations of the other compounds also increased or decreased, but the changes were not significant. The results showed that CtUGT3 exhibits 3-*O*-glycosylation in safflower and is involved in astragalin biosynthesis.

**Figure 5.**
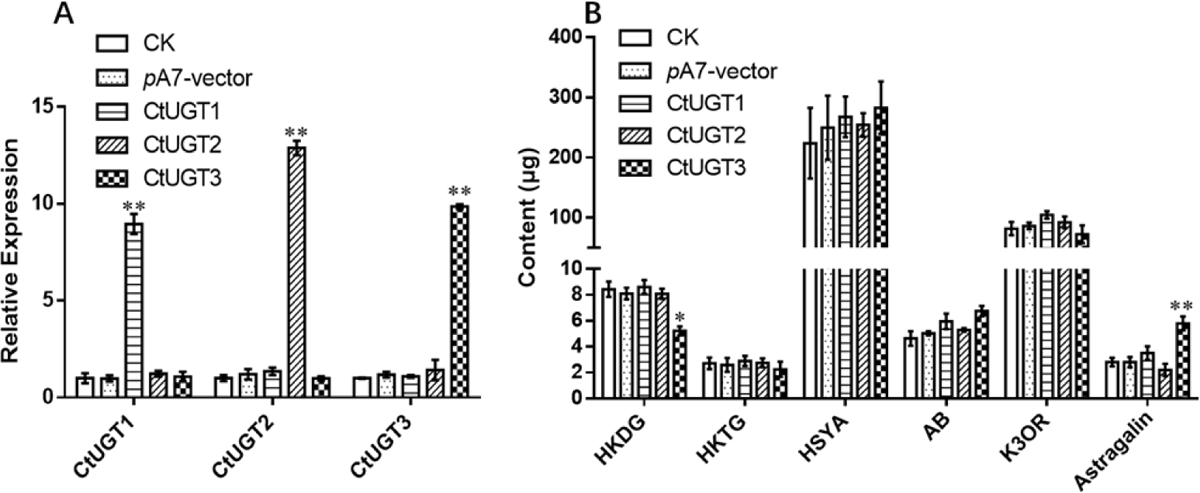
The relative expressions of *UGT*s and the contents of 6 flavonoid glycosides in corolla protoplast after transient expression of CtUGTs. (A) represents the relative expressions of *CtUGT1*, *CtUGT2* and *CtUGT3* in 5 different groups. (B) represents 6 flavonoid glycosides in corolla protoplast. HKDG (6-hydroxyl kaempferol 3,6-*O*-diglucosyl-7-*O*-glucuronic acid); HKTG (6-hydroxyl kaempferol 3,6,7-*O*-triglucoside); HSYA (hydroxysafflor yellow A); AB (anhydrosafflor yellow B); K3OR (kaempferol 3-*O*-rutinoside); Astragalin (kaempferol 3-*O*-glucoside). CK represents the untransfected group. *p*A7-vector represents the group of transfected empty vector. CtUGT1, CtUGT2 and CtUGT3 represent corolla protoplast samples that transient expressed *CtUGT1*, *CtUGT2* and *CtUGT3*, respectively. Some of the differences were marked in the figure (\**P* < 0.05, \*\**P* < 0.01).

### Verification of enzyme activity

The three recombinant UGT proteins were successfully expressed in the BL21 (DE3) strain and purified using the His-tag (Supplemental Figure S8). Flavonoid aglycones commonly present in safflower flowers were used as sugar acceptors to detect enzyme activities, such as kaempferol, apigenin, quercetin and isorhamnetin. Naringenin and naringenin chalcone were also selected as sugar receptors, because they are key nodes in the flavonoid biosynthesis pathway and safflower contains a large amount of chalcones. UDP-glucose was used as a sugar donor. Naringenin chalcone 2’-*O*-glucoside was elucidated by its ^1^H NMR spectrum (Supplemental Figure S9) and ^13^C NMR spectrum (Supplemental Figure S10), and the other products were matched with commercial standards. CtUGT3 catalyzed all the substrates and exhibited stronger glycosylation activity than the other two candidates.

When naringenin (**1**) was used as the acceptor substrate, CtUGT3 catalyzed the formation of naringenin 7-*O*-glucoside (**1a**) (Supplemental Figure S11). When naringenin chalcone (**2**) was used as the acceptor substrate, CtUGT3 catalyzed the formation of two glycosylation products with the same molecular weight. One product was identified as naringenin 2’-*O*-glucoside (**2a**) and the other was matched to **1a** (Supplemental Figure S12). Naringin chalcone and naringin are a pair of isomers. Naringin chalcone can be easily closed-loop converted into naringin in an aqueous solution, and naringin can also be open-loop converted to naringin chalcone in a strongly alkali environment. Therefore, CtUGT3 produced two products with the same molecular weight when **2** was used as the acceptor. However, **1** and **2** exhibited distinct UV spectrum, and their glycosylation products exhibited the same UV absorption. Compounds **1** and **1a** showed the highest UV absorption around 280 nm, whereas **2** and **2a** showed strong UV absorption around 360 nm. This feature can simply distinguish chalcones from dihydroflavones. CtUGT3 produced apigenin 7-*O*-glucoside (**3a**) when apigenin (**3**) was used as the acceptor (Supplemental Figure S13). Multiple products were identified when kaempferol (**4**), quercetin (**5**) or isorhamnetin (**6**) were glycosylated by CtUGT3 respectively (Supplemental Figure S14-S16). The three products were identified as kaempferol 7-*O*-glucoside (**4a**), kaempferol 3-*O*-glucoside (**4b**), and kaempferol 3,7-di-*O*-glucoside (**4c**) when kaempferol used as substrate. Catalytic reactions of CtUGT3 with **4a** and **4b** were also investigated. However, the obvious peak of **4c** was only observed when **4a** was used as the substrate, and it was difficult to convert **4b** into **4c**. The conversion rates of different aglycones catalyzed by CtUGT3 are shown in Figure 6.

**Figure 6.**
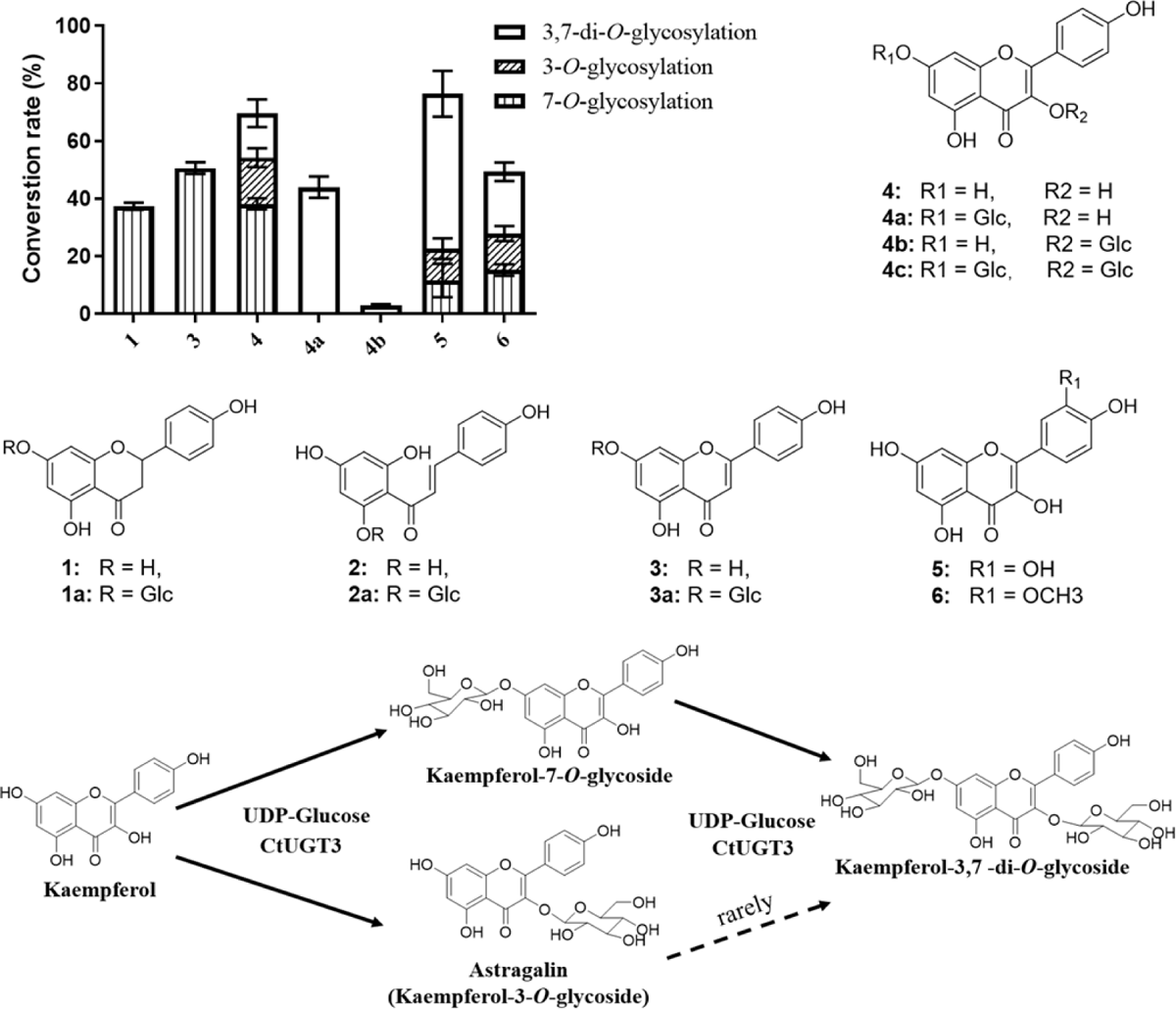
Conversion rates of flavonoid aglycones catalyzed by CtUGT3. Compounds 1, 3, 4, 4a, 4b, 5, and 6 represent naringenin, apigenin, kaempferol, kaempferol 7-*O*-glucoside, kaempferol 3-*O*-glucoside, quercetin, and isorhamnetin, respectively.

### Biochemical properties and kinetic parameter analysis for CtUGT3

To examine the biochemical properties of CtUGT3, the temperature, reaction time, divalent metal ions, and pH were optimized using **1** as the sugar acceptor and UDP-glucose as the sugar donor (Supplemental Figure S17). CtUGT3 exhibited the best activity in the range of 40 ∼ 50℃. The reaction was most intense in the first 5 min and was complete in approximately 60 min. In addition to the inhibitory effect of Co^2+^, most divalent metal ions had no obvious effect on the activity of CtUGT3. The activity of CtUGT3 was highest in Na_2_HPO_4_-NaH_2_PO_4_ buffer at pH 8.

The apparent *K*_m_ values were determined from Michaelis-Menten plots with varying concentrations of the sugar acceptors. When UDP-glucose was saturated, the apparent *K*_m_ values for **1**, **3**, **4, and 4a** were 70.16 μM, 75.46 μM, 110.40 μM, and 39.70 μM, respectively, and *K*_cat_/*K*_m_ values for **1**, **3**, **4, and 4a** were 999.2 M^-1^·s^-1^, 2207.5 M^-1^·s^-1^, 4116.8 M^-1^·s^-1^ and 2537.2 M^-1^·s^-1^, respectively (Supplemental Figure S18). Although the *K_m_* value reflected that the affinity between CtUGT3 and kaempferol was relatively weak, the efficiency of CtUGT3 catalyzing kaempferol was the highest as shown by the *K*_cat_/*K*_m_ value.

### Analysis of active site of CtUGT3 based on autodocking and mutagenesis

Eight amino acids, Asn244, Lys246, Ser276, His360, Asn364, Glu368, Glu384, and Gln385, showed interactions with kaempferol in autodocking results (Figure 7A). The three candidate CtUGTs and closely evolved UGTs were aligned. A reported 3-*O*-GT, Sb3GT1, was also compared with candidates, as CtUGT3 showed 3-*O*-glycosylated ability *in vivo*. Six amino acids were highly conserved among UGTs, including Ser276, His360, Asn364, Glu368, Glu384, and Gln385 (Supplemental Figure S19). To explore the key residues of CtUGT3 glycosylation, six mutants were cloned and used to catalyze kaempferol, including N244K, N244Q, K246Y, K246G, S276T, and E384D. In particular, E384D and S276T showed higher 3-*O*-glycoside content than the wild type (Figure 7B). Thus, kaempferol 7-*O*-glucoside was used as a substrate to further confirm the 3-*O*-glycosylation abilities of CtUGT3 and its mutants (Figure 7C). These results confirmed that the 3-*O*-glycoxylation abilities of E384D and S276T were superior than the others, and E384 and S276 were important amino acids for glycosylation activity of CtUGT3.

**Figure 7.**
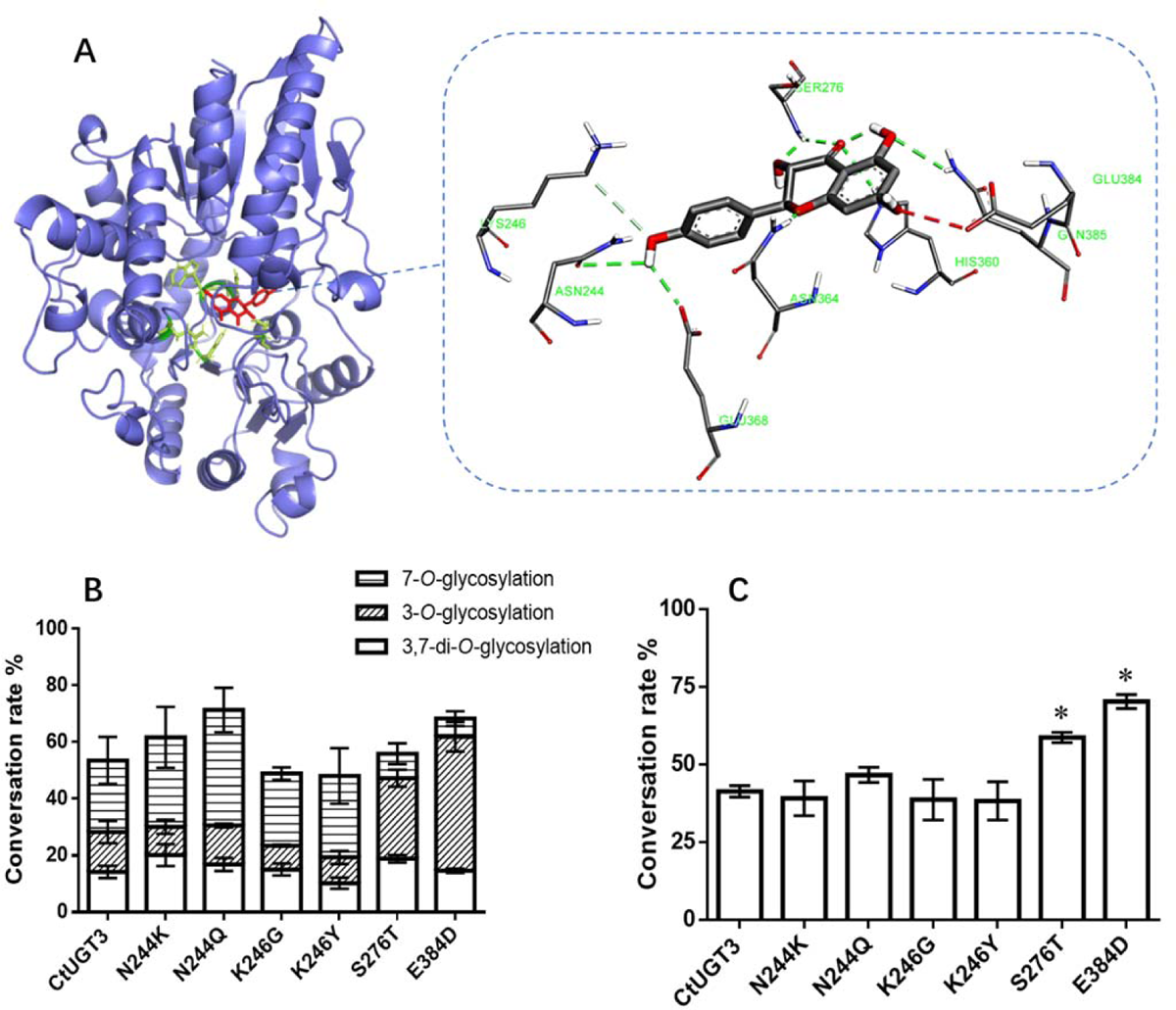
Molecular modeling and conversion rates of kaempferol and its 7-*O*-glucoside catalyzed by CtUGT3 and mutants. (A) model of CtUGT3 binding with kaempferol; (B) conversion rates of kaempferol catalyzed by CtUGT3 and mutants; (C) conversion rates of kaempferol 7-*O*-glucoside catalyzed by CtUGT3 and mutants. The differences were marked in the figure (\**P* < 0.05).

## Discussion

In this study, metabolomic and transcriptomic analyses were performed using safflower under different light quality treatments. The contents of the 9 flavonoid *O*-glycosides were significantly correlated with the expression of *CtUGT1*, *CtUGT2*, and *CtUGT3*. All these were 3-*O* and 7-*O* glycosides, like astragalin (kaempferol 3-*O*-glucoside) and kaempferol 7-*O*-glucoside. It has been speculated that CtUGTs may have 3-OH or 7-OH glycosylation abilities. We further confirmed that only CtUGT3 was involved in astragalin biosynthesis *in vivo* and exhibited 3-OH and 7-OH glycosylation activities *in vitro*.

Safflower flowers contain special flavonoids, such as chalcone *C*-glycosides, which are represented by HSYA and are the main components affecting safflower flower color rather than anthocyanins (Wang et al. 2021). However, model plants commonly used for functional identification may not contain these components or their precursors. In this experiment, protoplasts were prepared from young safflower corolla. In the protoplast that was successfully transformed into CtUGT3, obvious red fluorescence under DsRed was observed, and the cell color was also deeper. At the same time, the most common chalcones, kaempferol and its glycosides in safflower, such as HSYA, AB and astragalin, were identified in protoplasts, and the astragalin (kaempferol 3-*O*-glucoside) content was significantly increased. However, only kaempferol glycosides were detected in the protoplasts, and no glycosides of other flavonols such as quercetin and isorhamnetin were identified. It may be due to the fact that kaempferol is the most abundant flavonol in safflower (Xian et al. 2022), and the protoplasts were prepared from fresh, unflowering corollas, while metabolomics measured blooming flowers, resulting in slight differences in their compositions. On the other hand, although CtUGT3 exhibited 3-OH and 7-OH glycosylation activity *in vitro*, no significant changes were detected in any component containing 7-*O*-glycoside in corolla protoplasts. The results proved that CtUGT3 exhibited flavonoid *O*-glycosylation activities, especially kaempferol 3-OH glycosylation ability, in safflower flowers. The results also showed that the safflower corolla protoplast is an ideal material for verifying the GTs involved in flavonoid biosynthesis in safflower *in vivo*.

Glycosylation contributes to the diversity and complexity of plant secondary metabolites biosynthesis, and plays an important role in plant defense and stress resistance (Vogt and Jones 2000). Most flavonoids in plants exist as their glycosides, most of which are *O*-glycosides or *C*-glycosides. The most abundant flavonoid glycosides in plants are flavone *O*- or *C*-glycosides and flavonol *O*-glycosides, with *O*-glycosides at position 3 or 7 being the most common (Xiao 2017). Glycosyltransferases that catalyze flavonoid glycosides in plants have been reported a lot, especially 3-*O*-GT and 7-*O*-GT. Flavonoid 3-*O*-GT can glycosylate flavonols and anthocyanin aglycones. Sb3GT1 from *Scutellaria baicalensis* could 3-*O* glycosylate 17 flavonol aglycones, accepting five different sugar donors (UDP-Glc/-Gal/-GlcNAc/-Xyl/-Ara) (Wang et al. 2019). NpUGT6 has been identified in the embryos of *Nelumbo nucifera* and catalyzes the 3-OH glycosylation of quercetin (Hu et al. 2019). The expression of Fh3GT1 was positively correlated with flavonol glycoside and anthocyanin accumulation in *Freesia hybrida*. Transfection of Fh3GT1 into the *Arabidopsis* UGT78D2 mutant successfully restored the content of anthocyanins and flavonols caused by a loss in 3-*O*-GT function, indicating that Fh3GT1 plays the role of a flavonoid 3-*O*-GT (Sun et al. 2016). 7-*O*-GT is another common enzyme that catalyzes flavonoid glycosylation. A 7-*O*-GT from *Iris tectorum*, UGT73CD1, could glycosylate 7-OH and showed strong catalytic promiscuity for 12 types of flavonoids and 3,4-dichloroaniline to form *O*- or *N*-glycosides (Huang et al. 2021). A 7-*O*-GT, CsUGT75L12, has been identified in tea trees. Recombinant CsUGT75L12 showed glycosyltransferase activity at the 7-OH position of various flavonoids. Compared with the wild type, the flavonol glycoside content in *Arabidopsis* seeds overexpressing CsUGT75L12 was significantly higher (Dai et al. 2017).

Most GTs have broad substrate promiscuity, and some glycosylate at different positions for different substrate types. An *O*-GT in *Tripterygium wilfordii*, TwUGT2, has glycosyltransferase activity toward the 3-OH and 7-OH groups of flavonoids, thereby converting quercetin and pinocembrin into quercetin 3-*O*-glucoside and pinocembrin 7-*O*-glucoside, respectively (Lu et al. 2020). Another *O*-GT in *Tripterygium wilfordii*, TwUGT3, accepts luteolin, quercetin, pinocembrin, and genistein to produce luteolin 7-*O*-glucoside, quercetin 3-*O*-glucoside, pinocembrin 7-*O*-glucoside and genistein 4’-*O*-glucoside respectively (Gao et al. 2020). Forty-three *O*-GT candidate genes were screened from *Glycyrrhiza uralensis*. Eleven of candidates were characterized, including isoflavone 7-*O*-GTs, flavonol 3-*O*-GTs, and promiscuous *O*-GTs (Chen et al. 2019). In the present study, CtUGT3 showed 7-OH and 3-OH glycosylation activities at the same time when kaempferol was used as an aglycone *in vitro* and produced 3-*O*-glycoside, 7-*O*-glycoside, and 3,7-di-*O*-glycoside. It seems that CtUGT3 has stronger 7-OH catalytic activity than 3-OH glycosylation based on enzyme activity assay *in vitro*. However, when CtUGT3 was overexpressed in corolla cells, only a significant increase in astragalin (kaempferol 3-*O*-glycoside) was detected, instead of an increase in the content of 7-*O*-glycoside or 3,7-di-*O*-glucoside. On the other hand, although the *K_m_* value reflected that the affinity between CtUGT3 and kaempferol was relatively weak, the efficiency of CtUGT3 catalyzing kaempferol was the highest as shown by the *K*_cat_/*K*_m_ value. It was speculated that CtUGT3 might compete with or cooperate with other enzymes *in vivo*, which is different from the results of enzyme activity determination *in vitro*.

In safflower, three *UGT* genes related to flavonoid metabolism were screened through the safflower transcriptome, and their expression patterns were analyzed (Guo et al. 2016). UGT73AE1 was the first UGT identified in safflower. It is also the first reported trifunctional plant glycosyltransferase that exhibited the capability to glycosylate diverse acceptors to generate *N*-, *O*-, or S-glycosides (Xie et al. 2014). Subsequently, UGT71E5 was found in safflower (Xie et al. 2017). It can generate *N*-glycoside from the abundant *O*-glycoside and glycosylate 4 flavonoids (quercetin, naringenin, genistein and phloretin) to di-*O* or multi-*O*-glucosides. In this study, 3 *UGT* genes, which exhibited different flavonoid glycosylation abilities, were screened based on the metabolism and transcriptome of safflowers treated with different light qualities. Among these, CtUGT3 showed the strongest flavonoid glycosylation activity, both *in vivo* and *in vitro*. Our results confirmed that CtUGT3 could glycosylate flavonoid 7-OH and 3-OH, and exhibited 3-*O*-GT activity in astragalin biosynthesis in safflower.

To further understand CtUGT3 catalytic mechanism, autodocking was performed, which showed that 8 residues interacted with aglycone. In the docking results, S276 formed a hydrogen bond with the 3-OH group of kaempferol, and E384D formed a bond with 7-OH. However, both E384D and S276T mutants synthesized more 3-OH glycosylation products than the other mutants. D390 in GgCGT aligned with E384 in CtUGT3. Researchers have confirmed that sugar donor selectivity is controlled by hydrogen-bond interactions of sugar hydroxyl groups with D390 (Zhang et al. 2020). They found that D390E retained the same di-*C*-glycosylation activity as the wild-type, but D390A and D390N almost abolished *C*-glycosylation activity. In contrast, the E384D mutant showed increased 3-OH glycosylation ability of CtUGT3. Sb3GT1 is a 3-*O*-glycosyltransferase found in *Scutellaria baicalensis*. T281 in Sb3GT1 aligned with S276 in CtUGT3. T281 was thought to stabilize the sugar donor and the glycosylation function of T281A was almost lost (Wang et al. 2019). In our study, S276T exhibited stronger 3-OH glycosylation activity than the wild-type. Autodocking results showed that E384 interacted with 7-OH and S276 interacted with 3-OH (Figure 7A). Therefore, we speculated that 7-OH glycosylation might compete with 3-OH glycosylation when CtUGT3 catalyzed kaempferol, resulting in E384D mutant affecting 3-OH glycosylation activity and S276T obtaining stronger 3-OH glycosylation activity. E384 and S276 are critical catalytic residues for 3-OH glycosylation of CtUGT3.

## Conclusion

To reveal the UGTs involved in flavonoid glycoside biosynthesis in safflower flowers, metabolomic and transcriptome analyses were performed using safflower under different light quality treatments. Three differentially expressed *UGT* genes related to flavonoid 3-*O* or 7-*O*-glucosides (like kaempferol 3-*O*-glucoside and kaempferol 7-*O*-glucoside) contents were screened and transfected into corolla protoplasts. The astragalin (kaempferol 3-*O*-glucoside) content was more than doubled when CtUGT3 was overexpressed. The biochemical properties and kinetic parameters of CtUGT3 were also determined. It exhibited flavonoid 7-OH and 3-OH glycosylation activities *in vitro*. Molecular modeling and site-directed mutagenesis revealed that E384 and S276 were critical catalytic residues for 3-OH glycosylation of CtUGT3. These results demonstrate that CtUGT3 has a flavonoid 3-OH glycosylation function and is involved in the biosynthesis of astragalin in safflower. This study provides insights into the catalytic mechanisms of flavonoid *O*-glycosyltransferases, and makes a reference for flavonoid biosynthesis genes research in medicinal plants.

## Materials and methods

### Plant materials

Safflower flowers were cultivated in a phytotron. Prior to the experiment, the plants were grown at 25℃ for 12 h per day under 15,000 lux white light and at 18℃ in the dark; the relative humidity was set to 65%. Safflower plants of the same growth status were selected and grown in a peat: vermiculite: perlite (3:1:1) mixture. Before flowering, plants were divided into 6 groups of equally sized buds. The flowering process lasted for two days. During flowering, the 5 groups were illuminated for 12 h per day under different single light that emitted red light (664 nm), far-red light (724 nm), blue light (447 nm), UV-B light (290 nm), and white light, respectively. The last group was treated in dark. Then, blooming flowers were collected and stored at −80℃. Except for the far-red group, three biological replicates were used for the metabolomic and transcriptome analyses, and each replicate was mixed with 4 individuals. Due to the small amount of corollas in the far-red group, 12 individuals were mixed and detected three times.

### Liquid chromatography and mass spectrometry

Freeze-dried flowers were crushed using a mixer mill (MM 400, Retsch) with zirconia beads for 1.5 min at 30 Hz. One hundred milligrams of powder was weighted and extracted overnight at 4 ℃ with 0.6 mL of 70% aqueous methanol. Following centrifugation at 10, 000 × g for 10 min, the extracts were filtered and analyzed using a UPLC-ESI-MS/MS system (UPLC, Shim-pack UFLC SHIMADZU CBM30A system; MS, Applied Biosystems 4500 Q TRAP). The analytical conditions were as follows: UPLC: column, Waters ACQUITY UPLC HSS T3 C18 (1.8 μm, 2.1 mm*100 mm). The mobile phase comprised solvent A (pure water with 0.04% acetic acid) and solvent B (acetonitrile with 0.04% acetic acid). Sample measurements were performed using a gradient program with starting conditions of 95% A and 5% B. Within 10 min, a linear gradient of 5% A and 95% B was programmed, and the composition of 5% A and 95% B was maintained for 1 min. Subsequently, the composition was adjusted to 95% A and 5.0% B within 0.10 min and kept for 2.9 min. The column oven was set to 40°C, and the injection volume was 4 μL. The effluent was alternately connected to an ESI-triple quadrupole-linear ion trap (QTRAP)-MS. LIT and triple quadrupole (QQQ) scans were acquired on a triple quadrupole-linear ion trap mass spectrometer (Q TRAP), API 4500 Q TRAP UPLC/MS/MS System, equipped with an ESI Turbo Ion-Spray interface, operating in positive and negative ion modes and controlled by Analyst 1.6.3 software (AB Sciex). The ESI source operation parameters were as follows: ion source, turbo spray; source temperature 550℃; ion spray voltage (IS) 5500 V (positive ion mode)/-4500 V (negative ion mode); ion source gas I (GSI), gas II, (GSII) and curtain gas (CUR) were set at 50, 60, and 30.0 psi, respectively; and the collision gas (CAD) was high. Instrument tuning and mass calibration were performed with 10 and 100 μmol/L polypropylene glycol solutions in QQQ and LIT modes, respectively. QQQ scans were acquired as MRM experiments, with the collision gas (nitrogen) set to 5 psi. The DP and CE for individual MRM transitions were performed with further DP and CE optimization. A specific set of MRM transitions was monitored for each period according to the metabolites eluted within this period.

Metabolite identification and quantification were performed as previously described (Chen et al. 2013). Metabolite identification was based on the primary and secondary spectral data annotated by public databases, namely, MassBank (http://www.massbank.jp/), KNAPSAcK (http://kanaya.naist.jp/KNApSAcK/), HMDB (http://www.hmdb.ca/), MoToDB (http://www.ab.wur.nl/moto/), and METLIN (http://metlin.scripps.edu/index.php). Metabolite quantification was performed using MRM. The full dataset of the annotated metabolites is present in Supplemental Table S8.

Significantly regulated metabolites between groups were determined by VIP (variable importance in project) ≥ 1 and absolute Log2FC (fold change) ≥ 1. VIP values were extracted from the orthogonal partial least squares-discriminant analysis (OPLS-DA) results, which also contained score plots and permutation plots, and were generated using the R package MetaboAnalystR. The data were log transformed (log2) and mean centered prior to OPLS-DA. A permutation test (200 permutations) was performed to avoid overfitting.

The identified metabolites were annotated using the Kyoto Encyclopedia of Genes and Genomes (KEGG) compound database (http://www.kegg.jp/kegg/compound/), and the annotated metabolites were mapped onto the KEGG pathway database (http://www.kegg.jp/kegg/pathway.html). Pathways with mapped significantly regulated metabolites were then fed into the metabolite set enrichment analysis (MSEA), and their significance was determined using a hypergeometric test’s *p*-values.

### RNA sequencing and annotation

All samples were ground in liquid nitrogen, and total RNA was isolated by TRIzol reagent (Invitrogen, CA, USA). To remove DNA, an aliquot of total RNA was treated with DNase (Takara, Dalian, China) using the standard protocol described by the manufacturer. The purity of the RNA samples according to the A260/A280 ratio was determined using a NanoPhotometer; the A260/A280 ratios of all samples were in the approximate range of 1.9-2.1. The integrity of RNA samples was assessed using an Agilent 2100 Bioanalyzer, and samples with no signs of degradation were selected for further analysis.

Sixteen samples, including one mixed sample from the far-red group and three bioreplicates per group from the other five treatments, were used to construct the libraries. mRNA was isolated from total RNA using magnetic beads with oligo (dT) primers, cDNA was synthesized using a cDNA synthesis kit (TaKaRa, Dalian, China), and the sequencing adapter was linked to both ends. Library preparations were sequenced on an Illumina HiSeq 4000 platform. After quality control was performed, clean reads were compared to the reference genome (not published). The unigenes were annotated in the following databases: KEGG, NR, SwissProt, GO, KOG, and Trembl. The amino acid sequences of the Unigenes were aligned with the Pfam database by HMMER software. The raw data were submitted to NCBI (PRJNA831043).

### Analysis of DEGs

Clean reads were mapped to Unigenes by Bowtie2 v2.2.5 (Langmead and Salzberg 2012), and gene expression levels were calculated through RSEM v1.2.12 (Li and Dewey 2011). Differential expression analysis was performed by DEseq2 (Love et al. 2014; Varet et al. 2016). A fold change ≥ 2.00 and an adjusted *P* value ≤ 0.05 were set as the thresholds to establish significantly different expression. All DEGs were analyzed using KEGG and Gene Ontology (GO) enrichment.

### Real-time qPCR analysis

RT-qPCR was performed as previously described (Ren et al. 2020a). Each sample was replicated 3 times. Specific primers were designed by Primer Premier 5 software (Supplemental Table S9). Gene expression levels under different treatments were measured with the CFX96^TM^ Real-time System (Bio-Rad, USA) using SYBR Premix Ex Taq Ⅱ (TaKaRa, Japan). The *25S* rRNA gene obtained from *Carthamus tinctorius* L. was used as an internal reference gene to identify the differences in each cDNA template. The RT-qPCR cycling conditions were as follows: 95°C for 3 min, followed by 40 cycles of 95°C for 10 s and 61°C for 30 s. The 2^-ΔΔCT^ method (Livak and Schmittgen 2001) was employed to analyze relative gene expressions. The analysis of variance (ANOVA)-Tukey’s test in SPSS (version 20) was used for data analysis. Two PAL genes, *HCT* gene, CHS gene, *FLS* gene and three *UGT* genes related to flavonoid biosynthesis, namely, *CtPAL1*, *CtPAL2*, *CtHCT*, *CtCHS*, *CtFLS*, *CtUGT1*, *CtUGT2* and CtUGT3, were successfully amplified and selected to confirm the results.

### Molecular cloning and phylogenetic analysis

A mixed cDNA library containing cDNA from the different light treatments was constructed. Total RNA was isolated using TRIzol, after which first-strand cDNA was synthesized from total RNA using a cDNA reverse transcription kit (TaKaRa, Japan). Full-length CDS sequences of the three candidate UGT genes were obtained from third-generation transcriptome sequencing data (Chen et al. 2018). Specific primer pairs were designed by Primer Premier 5.0 (Supplemental Table S9). PCR was performed as follows: initial denaturation at 95°C for 4 min; 34 cycles of 95°C for 30 s, 58°C for 30 s, and 72°C for 2 min; and a final extension at 72°C for 8 min. The PCR products were cloned into pMD19-T vectors (TaKaRa, Japan). Different UGT amino acid sequences were downloaded from NCBI. UGT phylogenetic tree was constructed using the Geneious Prime software.

### Isolation of corolla protoplast

Protoplast isolation was performed according to previously reported methods with some modifications (Chen et al. 2015; Ren et al. 2020b). The tender corolla from *Carthamus tinctorius* was cut into 0.15 mm pieces and transferred into a freshly prepared enzyme solution [1.5 % Cellulase RS (Yakult, Japan), 0.3 % Macerozyme R-10 (Yakult, Japan), 0.5 M mannitol, 10 mM MES, 10 mM CaCl_2_, and 0.1% bovine serum albumin (BSA)] in a 50 mL flask. The enzymes were thermally activated at 55°C for 10 min before the addition of CaCl_2_ and BSA. The tissues were incubated on an orbital shaker at 40 rpm at 25°C for 3 h. Then, an equal volume of W5 (154 mM NaCl, 125 mM CaCl_2_, 5 mM KCl and 4 mM MES pH 5.7) was added to the enzyme hydrolysis solution and vigorously shaken for 5 s to stop the hydrolysis. A 40 µm nylon filter was used for filtration, and the filtrate was horizontally centrifuged at 100 × g for 2 min. The supernatant was discarded, and the protoplasts were resuspended in 5 mL of cold W5 solution and horizontally centrifuged at 100 × g for 1 min. The supernatant was discarded, and the protoplasts were resuspended in 5 mL of cold W5 solution. The suspension was incubated on ice for 30 min. The cells were then settled at the bottom by gravity. The upper liquid was carefully discarded, and the protoplasts were adjusted to a density of 1 × 10^5^–1 × 10^6^/mL with MMG (15 mM MgCl_2_, 0.5 M mannitol and 4 mM MES at pH 5.7).

### Transfection and transient expression of CtUGTs in corolla protoplast

CtUGTs were cloned into the pA7-CaMV35S-YFP vector for transient expression analysis of safflower corolla protoplasts. The protoplast transfection procedure followed that described in a previous study with slight modifications (Zhang et al. 2011). Then, 10 ∼ 20 µg plasmid was added into 100 µL of protoplast suspension and mixed gently. A total of 110 µL PEG (40% PED4000, 0.2 M mannitol, 100 mM CaCl_2_) was then added and mixed gently. The mixture was incubated at 25°C for 15 min, and then 440 µL of W5 was added to stop the transfection. The mixture was then centrifuged horizontally at 100 × g for 2 min. The supernatant was discarded and the protoplasts were resuspended in 1 mL of MMG. The protoplasts were incubated in the dark for 12 h and then used for experimental analysis.

After collecting the cells, 20 mg protoplasts were weighed and extracted with 5 mL water at 60°C for 45 min. The extracts were detected by UPLC-QTOF-MS/MS and separated on an Eclipse Plus C18 column (150 mm×3.0 mm,1.8 μm, Agilent, USA) with acetonitrile and H_2_O containing 0.1% formic acid (v/v) as the mobile phase. A linear gradient HPLC elution program was used as follows: 0 min, 5% acetonitrile; 25 min, 95% acetonitrile. Extracted ion chromatograms (EICs) of flavonoid glycosides were generated for comparison with commercial standards. Protoplasts transfected with the empty pA7 vector were used as negative controls. The differentially metabolized flavonoid glycosides were screened.

### Expression and purification of recombinant proteins

The three *UGT* genes were cloned into the pLATE52 vector (Thermo Fisher, USA) for prokaryotic expression analysis. pLATE52-CtUGTs recombinant vectors were transformed into the BL21 (DE3) strain using the heat-shock method. BL21 cells were grown at 37℃ in 1 L of LB in a shake flask supplemented with 50 μg/mL ampicillin to an OD600 of 0.5 at 200 rpm. The flask was then placed on ice for 10 min and induced with 120 μM of IPTG. The cells were then incubated for 16 h at 16℃ and 200 rpm.

The cells were harvested by centrifugation (6517g, 15 min, 4℃), and the supernatant was removed. The cell pellet was resuspended in 30 mL of lysis buffer (25 mM HEPES pH 8, 500 mM NaCl, 5 mM imidazole), and the cells were lysed by sonication on ice. Cellular debris was removed by centrifugation (12,516 g, 45 min, 4℃), and the supernatant was filtered with a 0.45 μm filter before batch binding. Ni-NTA resin (Qiagen) was added to the filtrate at 1.5 mL/L of cell culture, and the samples were nutated for 1 h at 4℃. The protein–resin mixture was loaded onto a gravity flow column. The flow-through was discarded and the column was washed with approximately 30 mL of wash buffer (25 mM HEPES pH 8, 100 mM NaCl, 20 mM imidazole). The tagged protein was eluted in approximately 20 mL of elution buffer (25 mM HEPES pH 8, 100 mM NaCl, 250 mM imidazole). The entire process was monitored using Bradford assay. CtUGTs were concentrated using a 30 kDa Amicon Ultra spin filter, and 10% v/v glycerol was added. The proteins were flash-frozen in liquid nitrogen and stored at −80℃.

### Enzyme activity assay

In a total volume of 100 μL, 0.5 mM uridine diphosphate glucose (UDP-Glc) and 0.2 mM naringenin (**1**), naringenin chalcone (**2**), apigenin (**3**), kaempferol (**4**), quercetin (**5**) and isorhamnetin (**6**) were incubated with a 50 mM NaH_2_PO_4_-Na_2_HPO_4_ buffer (pH 8.0) containing 10 μM of purified CtUGTs at 30°C for 30 min. Heat-inactivated proteins (boiling at 100 °C for 10 min) were used for the negative control reaction. The reactions were quenched with 200 μL ice cold methanol (MeOH) and centrifuged at 12,000 rpm for 20 min. The supernatants were analyzed by UPLC-QTOF-MS/MS, as described above. The glycosylation products were identified by NMR or were matched with commercial standards. The percent conversion rate reflected how many substrates were converted into products. The percent conversion rates were calculated from the peak areas using standard curves of the glycosylated products and substrates.

### Determination of the biochemical properties and kinetic parameters of CtUGT3

The biochemical properties of CtUGT3 were investigated using 1 mM naringenin (**1**) as the sugar acceptor and 2 mM UDP-Glc as the sugar donor in 100 μL reaction mixtures that included 10 μM of purified protein. To determine the optimal reaction time for CtUGT3, assays were performed with different reaction time, including 1, 2, 5, 10, 20, 30, 60, and 120 min. To determine the optimal pH value for CtUGT3 activity, assays were performed in different buffers with pH ranges of 4.0 ∼ 6.0 (50 mM citric acid-sodium citrate buffer), 6.0 ∼ 8.0 (50 mM Na_2_HPO_4_-NaH_2_PO_4_ buffer), 7.0 ∼ 9.0 (50 mM Tris-HCl buffer), and 9.0 ∼ 11.0 (50 mM Na_2_CO_3_-NaHCO_3_ buffer).

The system was incubated at different temperatures (4, 16, 25, 37, 50, 60, and 70°C) to optimize the reaction temperature for CtUGT3 activity. To determine the dependence of divalent metal ions on CtUGT3 activities, Ca^2+^, Co^2+^, Fe^2+^, Mg^2+^, Mn^2+^, Zn^2+^, and EDTA were used individually at a final concentration of 5 mM. Three parallel reactions were performed for each condition. The reactions were quenched with ice-cold methanol and centrifuged at 12,000 rpm for 20 min. The supernatants were analyzed by HPLC as described above.

To determine the kinetic parameters of CtUGT3 for naringenin, assays containing 50 mM Na_2_HPO_4_-NaH_2_PO_4_ (pH 8.0), 1 μM CtUGT3, saturating UDP-glucose, and varying concentrations (5, 10, 20, 40, 80, 100, 200, 400, 800, 1000 and 1500 µM) of naringenin, were performed at 37 °C for 5 min with a final volume of 100 μL. To determine the kinetic parameters of CtUGT3 for apigenin, assays containing 50 mM Na_2_HPO_4_-NaH_2_PO_4_ (pH 8.0), 1 μM CtUGT3, saturating UDP-glucose, and varying concentrations (5, 10, 20, 40, 80, 100, 200, 400, 800, 1000 and 1500 µM) of apigenin, were performed at 37 °C for 5 min with a final volume of 100 μL. To determine the kinetic parameters of CtUGT3 for kaempferol, assays containing 50 mM Na_2_HPO_4_-NaH_2_PO_4_ (pH 8.0), 1 μM CtUGT3, saturating UDP-glucose, and varying concentrations (100, 200, 400, 800, 1000, 1200, 1600, 1800, 2000 and 2400 µM) of kaempferol, were performed at 37 °C for 5 min with a final volume of 100 μL. To determine the kinetic parameters of CtUGT3 for kaempferol 7-*O*-glycoside, assays containing 50 mM Na_2_HPO_4_-NaH_2_PO_4_ (pH 8.0), 1 μM CtUGT3, saturating UDP-glucose, and varying concentrations (5, 10, 20, 40, 80, 100, 200, 400, 600, 800, and 1200 µM) of kaempferol 7-*O*-glycoside, were performed at 37 °C for 5 min with a final volume of 100 μL. To determine the kinetic parameters for the UDP-glucose assays containing 50 mM Na_2_HPO_4_-NaH_2_PO_4_ (pH 8.0), 1 μM CtUGT3, saturating naringenin, and varying concentrations (1, 2.5, 5, 10, 20, 40, 80, 100, 200 and 400 µM) of UDP-glucose, were performed at 37°C for 5 min with a final volume of 100 μL. All reactions were quenched with ice-cold MeOH and centrifuged at 12,000 rpm for 10 min, and the supernatants were analyzed by HPLC. All experiments were performed in triplicates. The value of *K_m_* was calculated using a Michaelis-Menten plot.

### Molecular docking and mutagenesis

The protein models of CtUGT3 were established by SWISS-MODEL. Molecular docking of CtUGT3 with kaempferol was performed by AutoDock Tools (Trott and Olson 2010). The complex structure with the lowest binding energy was used in further studies. The desired mutants were constructed by site-directed mutagenesis of CtUGT3 using the wild-type expression plasmid as the template for PCR. PrimeSTAR Max Premix (2x) (TaKaRa, Japan) was used for PCR with the degenerate primers listed in Supplemental Table S10. The exact PCR conditions were as follows: 95 ℃ for 1 min, followed by 30 cycles of 95 ℃ for 10 s, 61 ℃ for 5 s, and 72 ℃ for 10 s, end with 72 ℃ for 1 min. The mutants, verified by sequencing, were transformed into *E.coli* BL21 (DE3) for the heterologous expression and purification of recombinant proteins as described above. Enzyme activity assays of the CtUGT3 mutants were performed with the same method used for the wild-type enzyme. The percent conversion rates were calculated from the peak areas using standard curves of the glycosylated products and substrates.

## Acknowledgements

This project is supported by grants from the National Natural Science Foundation of China (82274039, U19A2010), Department of Science and Technology of Sichuan Province (2021YFYZ0012-5, 2020YFN0152), Natural Science Foundation of Sichuan Province (2023NSFSC1770), State Administration of Traditional Chinese Medicine of the People’s Republic of China (ZYYCXTD-D-202209), and Chengdu University of Traditional Chinese Medicine “Xinglin Scholar” Research Enhancement Program (BSH2023014). We would like to thank Editage (www.editage.cn) for English language editing (JOB CODE: JIUAJ_6).

## Author contributions

CR performed the experiments, analyzed the data and wrote the manuscript. ZX helped with enzyme activity analysis. BX helped with genes cloning and expressions analysis. CC helped with protoplasts preparation. XH and HJ helped with autodocking analysis. JP and CP conceived and designed the study. JP and JC supervised the experiments.

## Supporting Information

Additional Supporting Information may be found in the online version of this article.

**Figure S1.** FPKM of 8 differential expressed genes related with flavonoid biosynthesis.

**Figure S2.** LC/MS analyses of 6-hydroxyl kaempferol 3,6-*O*-diglucosyl-7-*O*-glucuronic acid in protoplast. (A) Extracted ion chromatogram (EIC) of 801 in protoplast sample, and its MS and MS/MS spectra. (B) Total ion chromatogram of 6-hydroxyl kaempferol 3,6-*O*-diglucosyl-7-*O*-glucuronic acid (HKDG) standard, and its MS and MS/MS spectra.

**Figure S3.** LC/MS analyses of 6-hydroxyl kaempferol 3,6,7-*O*-triglucoside in protoplast. (A) EIC of 787 in protoplast sample, and its MS and MS/MS spectra. (B) Total ion chromatogram of 6-hydroxyl kaempferol 3,6,7-*O*-triglucoside (HKTG) standard, and its MS and MS/MS spectra.

**Figure S4.** LC/MS analyses of HSYA in protoplast. (A) EIC of 611 in protoplast sample, and its MS and MS/MS spectra. (B) Total ion chromatogram of HSYA standard, and its MS and MS/MS spectra.

**Figure S5.** LC/MS analyses of anhydrosafflor yellow B in protoplast. (A) EIC of 1043 in protoplast sample, and its MS and MS/MS spectra. (B) Total ion chromatogram of anhydrosafflor yellow B (AB) standard, and its MS and MS/MS spectra.

**Figure S6.** LC/MS analyses of kaempferol 3-*O*-rutinoside in protoplast. (A) EIC of 593 in protoplast sample, and its MS and MS/MS spectra. (B) Total ion chromatogram of kaempferol 3-*O*-rutinoside (K3OR) standard, and its MS and MS/MS spectra.

**Figure S7.** LC/MS analyses of astragalin in protoplast. (A) EIC of 447 in protoplast sample, and its MS and MS/MS spectra. (B) Total ion chromatogram of astragalin (kaempferol 3-*O*-glucoside) standard, and its MS and MS/MS spectra.

**Figure S8.** Protein purification of CtUGTs in safflower.

**Figure S9.** The ^1^H NMR spectrum of naringenin chalcone 2’-*O*-glucoside. Figure S10. The ^13^C NMR spectrum of naringenin chalcone 2’-*O*-glucoside. Figure S11. LC/MS analyses of CtUGTs enzyme activity assays with naringenin. (A-C) Total ion chromatogram (TIC) and EIC of naringenin and its glycosylated products in CtUGTs catalytic reactions. (D-E) MS spectra of naringenin and its 7-*O*-glucoside.

**Figure S12.** LC/MS analyses of CtUGTs enzyme activity assays with naringenin chalcone. (A-C) TIC and EIC of naringenin chalcone and its glycosylated products in CtUGTs catalytic reactions. (D-F) MS spectra of naringenin chalone and its glycosylated products.

**Figure S13.** LC/MS analyses of CtUGTs enzyme activity assays with apigenin. (A-C) TIC and EIC of apigenin and its glycosylated products in CtUGTs catalytic reactions. (D-E) MS spectra of apigenin and its glycosylated products.

**Figure S14. LC/MS analyses of CtUGTs enzyme activity assays with kaempferol.** (A-C) TIC and EIC of kaempferol and its glycosylated products in CtUGTs catalytic reactions. (D-G) MS spectra of kaempferol and its glycosylated products.

**Figure S15. LC/MS analyses of CtUGTs enzyme activity assays with quercetin.** (A-C) TIC and EIC of quercetin and its glycosylated products in CtUGTs catalytic reactions. (D-F) MS spectra of quercetin and its glycosylated products.

**Figure S16.** LC/MS analyses of CtUGTs enzyme activity assays with isorhamnetin. (A-C) TIC and EIC of isorhamnetin and its glycosylated products in CtUGTs catalytic reactions. (D-F) MS spectra of isorhamnetin and its glycosylated products.

**Figure S17.** Biochemical properties of CtUGT3. (A-D) Effects of temperature, reaction time, divalent metal ions and pH on CtUGT3 activity.

**Figure S18.** Determination of kinetic parameters for CtUGT3. (A-E) Determination of kinetic parameters of CtUGT3 for naringenin, apigenin, UDP-Glucose, kaempferol and kaempferol 7-*O*-glucoside, respectively.

**Figure S19.** Sequence alignment of CtUGT3 with its homologs.

**Table S1.** Flavonoids metabolites of safflower flowers treated by different light quality.

**Table S2.** Differentially metabolized flavonoids.

**Table S3.** Clean reads quality metrics.

**Table S4.** Statistic of Mapped Reads.

**Table S5.** Annotations of all DEGs.

**Table S6.** Statistic of differential expressed genes for different groups.

**Table S7.** Bioinformatics analysis of 3 candidate UGTs.

**Table S8.** Full dataset of annotated metabolites.

**Table S9.** Primers for RT-qPCR and cloning.

**Table S10.** Primers for CtUGT3 mutants cloning.

